# Resting-state brain activity can predict target-independent aptitude in fMRI-neurofeedback training

**DOI:** 10.1101/2021.02.08.430334

**Authors:** Takashi Nakano, Masahiro Takamura, Haruki Nishimura, Maro Machizawa, Naho Ichikawa, Atsuo Yoshino, Go Okada, Yasumasa Okamoto, Shigeto Yamawaki, Makiko Yamada, Tetsuya Suhara, Junichiro Yoshimoto

## Abstract

Neurofeedback (NF) aptitude, which refers to an individual’s ability to change its brain activity through NF training, has been reported to vary significantly from person to person. The prediction of individual NF aptitudes is critical in clinical NF applications. In the present study, we extracted the resting-state functional brain connectivity (FC) markers of NF aptitude independent of NF-targeting brain regions. We combined the data in fMRI-NF studies targeting four different brain regions at two independent sites (obtained from 59 healthy adults and six patients with major depressive disorder) to collect the resting-state fMRI data associated with aptitude scores in subsequent fMRI-NF training. We then trained the regression models to predict the individual NF aptitude scores from the resting-state fMRI data using a discovery dataset from one site and identified six resting-state FCs that predicted NF aptitude. Next we validated the prediction model using independent test data from another site. The result showed that the posterior cingulate cortex was the functional hub among the brain regions and formed predictive resting-state FCs, suggesting NF aptitude may be involved in the attentional mode-orientation modulation system’s characteristics in task-free resting-state brain activity.

## 1. Introduction

Neurofeedback (NF) with functional magnetic resonance imaging (fMRI-NF) is a non-invasive method that modulates brain activity patterns through procedural learning. It is expected to be a promising method for elucidating the mechanisms of brain functions and treating the brain abnormalities of several neuropsychiatric disorders (Sitaram et al., 2017; Stoeckel et al., 2014; Thibault et al., 2018). A growing body of literature has suggested the efficacy of fMRI-NF treatment for major depressive disorder (Mehler et al., 2018; Takamura et al., 2020; Young et al., 2017a, 2017b), bipolar disorder (Paret et al., 2016), schizophrenia (Ruiz et al., 2013; Zweerings et al., 2019), post-traumatic stress disorder (Gerin et al., 2016), attention-deficit hyperactive disorder (Alegria et al., 2017), anxiety disorder (Scheinost et al., 2013; Zilverstand et al., 2015), and Parkinson’s disease (Subramanian et al., 2011).

However, in terms of clinical usefulness, several issues remain with fMRI-NF, especially since an individual’s ability to change its brain activity through NF training, referred to as ‘NF aptitude’ in this paper, significantly varies across individuals (Kadosh and Staunton, 2019). The issue becomes more serious in the clinical application of fMRI-NF because individuals who have insufficient NF aptitude will barely benefit from fMRI-NF treatment. Changes in treatment are generally time-consuming and expensive. Therefore, when fMRI-NF is used for clinical practice, before such treatment, predicting whether each patient has sufficient NF aptitude is desirable.

The factors that define successful NF training can be broken down into individual participant factors as well as such external factors as experimental conditions (Haugg et al., 2020a; Sepulveda et al., 2016). Furthermore, individual aspects can be categorized into demographic-psychological (Kadosh and Staunton, 2019) and biological factors in which the neurobiological aspects are of our interest.

At present, the most extensive report on the predictions of individual NF aptitudes based on brain activity is the work by Haugg et al. (2020b), they gathered a large dataset from various types of fMRI-NF studies and tried to predict individual NF performances based on brain activation during localizer tasks in each experiment. More preferably, we wish to screen patients before NF training using a *task-free* setting, such as their resting-states. To the best of our knowledge, only one study predicted the efficacy of NF treatment/training based on resting-state FC (Scheinost et al., 2014). Unfortunately, it only focused on the NF performance of a particular target brain region and the resting-state FC connected to it. Considering that the target region of fMRI-NF treatment depends on the target disease or its symptoms, it is more desirable to predict individual aptitudes of NF targeting an arbitrary brain region based on the activity of widely-distributed brain regions, as we successfully predicted the individual characteristics related to cognitive functions and clinical states by combining the whole-brain FC with a machine learning technique (Yamashita et al., 2018; Yoshida et al., 2017).

Motivated by this analogy, in the present study, we examined the feasibility that an approach that combines the resting-state FC in the whole brain with a machine learning method can predict individual target-independent NF aptitudes.

## 2. Materials and Methods

### 2.1. Participants

For this study’s discovery dataset, 41 participants were recruited from the Hiroshima University Hospital, nearby medical clinics, or the local community. Among them, 35 participants had no history of mental or neurological disease. 22 participants had undergone neurofeedback training for left dorsolateral prefrontal cortex (DLPFC), and 13 for posterior cingulate cortex (PCC). The remaining six participants suffered from a major depressive disorder (MDD), had failed to respond to medication at least for four weeks, and underwent neurofeedback training for DLPFC.

For the test dataset, we recruited 24 healthy volunteers from the National Institute of Radiological Sciences, National Institutes for Quantum and Radiological Science and Technology. None had any history of neurological or psychiatric disorders. 16 participants had undergone neurofeedback training for pregenual anterior cingulate cortex (pgACC), and the remaining eight for intraparietal sulcus (IPS). All participants provided written informed consent before joining the study, which was approved by the Hiroshima University Clinical Research Review Committee or the National Institutes for Quantum and Radiological Science and Technology Certified Review Board in accordance with the ethical standards established by the 1964 Declaration of Helsinki and its later amendments. The details of the demographic statistics for both datasets are summarized in Table 1.

**Table 1:**
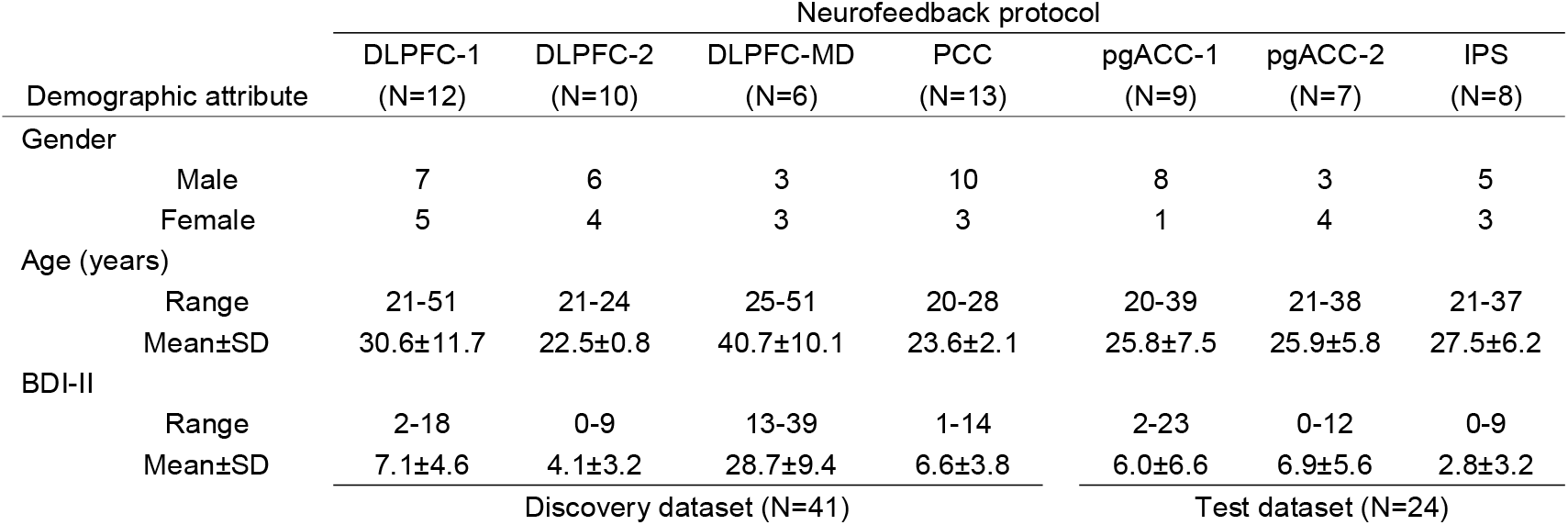
Demographics of participants in this study. BDI-II: Beck Depression Inventory (Second Edition).

All the participants completed the following identical procedure. After informed consent, they were administered BDI-II to evaluate their depressive symptoms outside of the MRI scanners. Next we acquired 10-minute (or 6-minute only for pgACC-1), eye-open resting-state fMRI, and structure MRI. During the resting-state fMRI scan, participants looked at a fixation point. The detailed scan parameters are displayed in Table 2. Following the resting fMRI session, participants underwent fMRI neurofeedback training. The procedures of the training session are described in the following section.

**Table 2:** Parameters of MRI scanners.

|  | Neurofeedback protocol |  |  |  |  |  |  |
| --- | --- | --- | --- | --- | --- | --- | --- |
|  | DLPFC-1 | DLPFC-2 | DLPFC-MD | PCC | pgACC-1 | pgACC-2 | IPS |
| MRI Scanner | Siemens MAGNETOM Verio 3.0T |  |  |  |  |  |  |
| resting state fMRI |  |  |  |  |  |  |  |
| scan time (mm:ss) | 10:13 | 10:20 | 10:13 | 10:13 | 6:52 | 10:13 | 10:13 |
| TR (ms) | 1000 | 2000 | 1000 | 1000 | 2000 | 1000 | 1000 |
| TE (ms) | 30 | 30 | 30 | 30 | 25 | 30 | 30 |
| flip angle (deg.) | 80 | 62 | 80 | 80 | 90 | 80 | 80 |
| voxel size (mm) | 3.3 x 3.3 x 3.2 | 2.0 x 2.0 x 2.0 | 3.3 x 3.3 x 3.2 | 3.3 x 3.3 x 3.2 | 3.8 x 3.8 x 3.8 | 3.3 x 3.3 x 3.2 | 3.3 x 3.3 x 3.2 |
| field of view (mm) | 212 | 192 | 212 | 212 | 240 | 212 | 212 |
| slice thickness (mm) | 3.2 | 2 | 3.2 | 3.2 | 3.8 | 3.2 | 3.2 |
| slice gap (mm) | 0.8 | 0 | 0.8 | 0.8 | 0.456 | 0.8 | 0.8 |
| slice number | 42 | 69 | 42 | 42 | 33 | 42 | 42 |
| T1w structure MRI |  |  |  |  |  |  |  |
| scan time (mm:ss) | 5:12 | 7:32 | 5:12 | 5:12 | 4:33 | 5:12 | 5:12 |
| TR (ms) | 2300 | 5000 | 2300 | 2300 | 2300 | 2300 | 2300 |
| TE (ms) | 2.98 | 2.88 | 2.98 | 2.98 | 1.98 | 2.98 | 2.98 |
| flip angle (deg.) | 9 | 4 | 9 | 9 | 9 | 9 | 9 |
| voxel size (mm) | 1.0 x 1.0 x 1.0 | 1.0 x 1.0 x 1.0 | 1.0 x 1.0 x 1.0 | 1.0 x 1.0 x 1.0 | 1.0 x 1.0 x 1.0 | 1.0 x 1.0 x 1.0 | 1.0 x 1.0 x 1.0 |
| field of view (mm) | 256 | 256 | 256 | 256 | 250 | 256 | 256 |
| slice thickness (mm) | 1 | 1 | 1 | 1 | 1 | 1 | 1 |
| slice number | 176 | 176 | 176 | 176 | 176 | 176 | 176 |
|  | Discovery dataset |  |  |  | Test dataset |  |  |

### 2.2. Neurofeedback training

#### 2.2.1. General NF protocol

In all datasets, we administered fMRI neurofeedback training to manipulate brain activity in a single region of interest (ROI) (Figure 1). On the feedback screen, the BOLD signal values for the participant ROIs were displayed as line graphs, alternating between blocks of up-regulation and down-regulation over the course of a single trial. The blocks were indicated by background color. The number of neurofeedback trials/visits and methods for defining the ROIs varied across the dataset. The following are the details of the procedures of each dataset.

**Figure 1:**
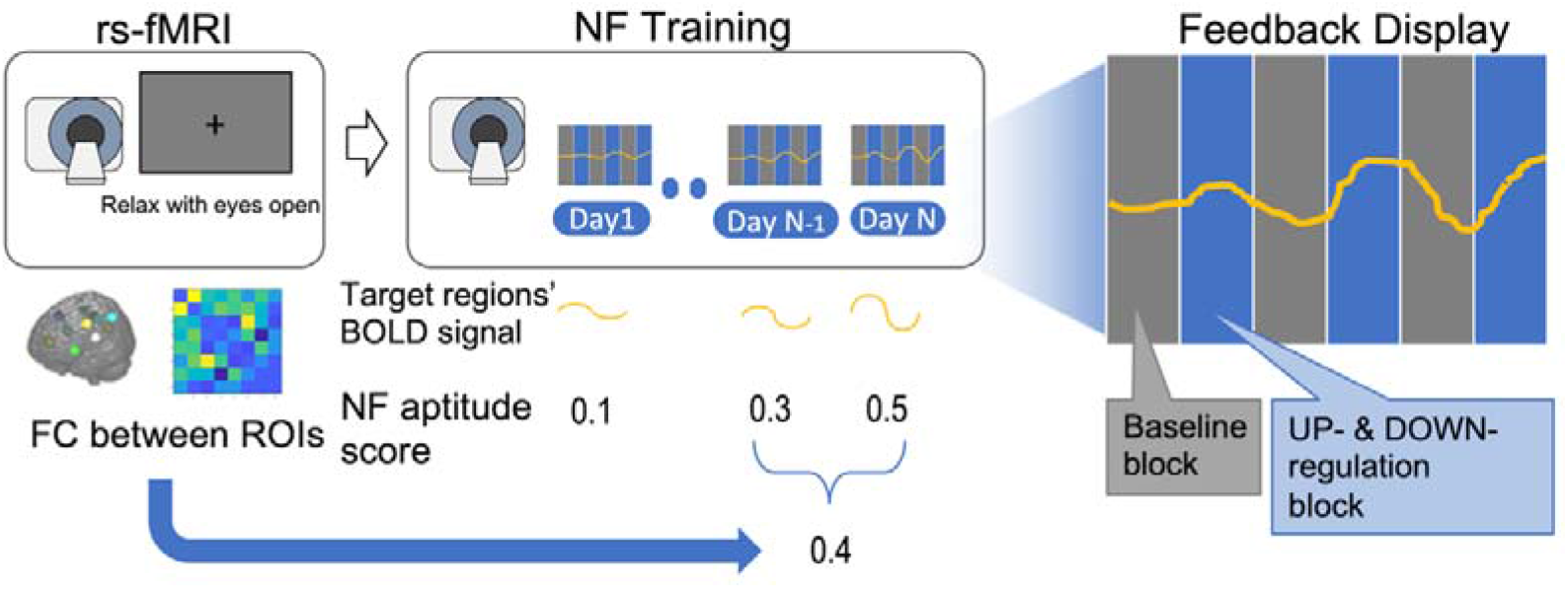
Diagram of data acquisition procedure. Participants underwent a resting-state fMRI (rs-fMRI) scan and then participated in the following neurofeedback (NF) training sessions. From the rs-fMRI data, a functional connectivity (FC) map, which is the synchronization between a set of brain regions, was calculated and used as a feature of the individual’s brain function. In the NF training sessions, participants were asked to increase or decrease their brain activity displayed on the feedback screen as a line graph. The degree of increase or decrease of the signal compared to baseline in each session was quantified as the NF aptitude scores. Finally, a model was created to predict the NF aptitude scores at the end of training from the FC map of the rs-fMRI sessions.

#### 2.2.2. Group-specific NF protocols

##### DLPFC-1

Each participant underwent 5-day neurofeedback training on an intermittent schedule over two weeks. On the first pre-training day, the resting-state fMRI data were obtained prior to collecting the first structural MRI scan. The ROI was defined as a 10-mm sphere at x = −44, y = 4, and z = 24 in the standard MNI brain. We used an EPI localizer scan to fit the ROI to the individual brain space of each participant. Neurofeedback training was composed of five baseline (rest) blocks (30 sec) and five up-regulation blocks (30 sec). They attempted several strategies (e.g., mental calculation, covert word operation) for increasing their neurofeedback signals during the up-regulation blocks. Real-time fMRI processing and feedback presentation were performed with Turbo-BrainVoyager™ version 3.2 (Brain Innovation, Maastricht, Netherlands).

##### DLPFC-2

Each participant underwent 3 days of neurofeedback training on an intermittent schedule over one week. At the start of each NF session, participants completed a 2-back task as a functional localizer scan (Sherwood et al., 2016) from which we generated participant-specific ROIs. The ROIs were defined as the following overlap region: 1) task-related activation in the localizer 2-back task, 2) voxels with negative FC with the PCC during the resting state, and 3) the anatomical left DLPFC region adapted from the Brainvisa Sulci atlas (http://brainvisa.info). NF training was composed of four repetitions of a baseline (rest) block (48 sec) and an up-regulation block (48 sec). Participants were instructed to raise their signals in the up-regulation block over their signals in the baseline block. Real-time fMRI processing and feedback presentation were performed with a Turbo-BrainVoyager™ version 3.2. Participants completed four NF runs during each session.

##### DLPFC-MD

The patients with MDD participated in successive 5-day neurofeedback training. The ROI-making and neurofeedback training procedures were identical as the DLPFC-2 group, except for the number of neurofeedback runs completed in each session. The patients were asked to complete 1 run; if they wished to continue, up to two additional runs were performed.

##### PCC

Each participant underwent 5-day neurofeedback training on an intermittent schedule over two weeks. On each day, resting-state fMRI data were obtained before and after the neurofeedback training (although the data were ignored in this study). The ROI was defined as a 10-mm sphere centered at MNI coordinates: x = −7, y = −49, and z = 28. We used an EPI localizer scan to fit the ROI to the individual brain spaces of each participant. Neurofeedback training was composed of five baseline blocks (30 sec) and five down-regulation blocks (30 sec). The participants tried several strategies (e.g., concentration on body sensations) to decrease their neurofeedback signals during the down-regulation blocks. Real-time fMRI processing and feedback presentation were performed with Turbo-BrainVoyager™ version 3.2 and MATLAB^®^ (Mathworks Inc., USA).

##### pgACC-1

Each participant underwent 4-day neurofeedback training on an intermittent schedule over two weeks. We used a functional localizer scan designed to efficiently locate in each participant the specific pgACC area that was most strongly involved in the processing of negative affective information. In the 7-min localizer scan, participants passively viewed happy or sad emotional faces. Following a localizer scan, they engaged in neurofeedback training, which was composed of five down-regulation blocks (32 sec) and five rest (baseline) blocks (32 sec). They tried several strategies to increase their positive mood and used them during the down-regulation blocks to increase the inverted neurofeedback signals. Online data analysis was performed using Turbo-Brain Voyager™ version 3.2.

##### pgACC-2 & IPS

Participants were randomly assigned to either the pgACC or IPS groups in a double-blinded manner such that the experimenters could not identify a targeted ROI for a given participant. Each participant underwent 4-day neurofeedback training on an intermittent schedule over two weeks. The neurofeedback training was composed of the following: (1) rest (30 sec ×1), followed by (2) up-regulation blocks (30 sec ×8) and (3) down-regulation blocks (30 sec ×8). Participants tried to increase and decrease their neural activities to match the inverted neurofeedback signals presented on the screen. Although experimenters did not suggest any strategies, participants were given hints that “positive” mood or thoughts may increase the signals, and “negative” moods or thoughts might decrease them. Previously described anatomical ROIs were specified as feedback targets: bilateral pgACC (7-mm radius sphere centered at MNI coordinates: x = 0, y = 46, and z = 4; (Ito et al., 2017)) and left IPS (7-mm radius sphere centered at MNI coordinates: x = −42, y = −51, and z = 52; (Young et al., 2017b)).

### 2.3 Neurofeedback aptitude scoring

To cope with the performance fluctuation over the sessions, we evaluated the individual NF aptitudes in the following procedures. For each session, let *BOLD_t_* be a BOLD signal averaged over the target ROI at the *t*-th time point. The signals were centered and normalized based on robust Z scores during the baseline periods. The normalized signals for the at the *t*-th time point were defined as

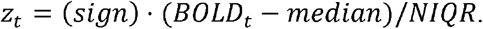

Here, *median* and *NIQR* are the median and the normalized interquartile range over the BOLD signals during the baseline period: {*BOLD_t_*: *t* ∈ (baseline periods)}; and (*sign*) is +1 if the condition of the neurofeedback training is for up-regulation or −1 for down-regulation. Using the normalized signals, the performance index for the *s*-th session on the *d*-th training day was evaluated as the average over the up-regulation (or down-regulation) periods:

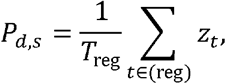

where (reg) denotes the set of the time points included in the up-regulation (or down-regulation) periods in the session and *T_reg_* is the total number of time points in (reg). Subsequently, the daily performance for the *d*-th day, denoted by 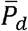 was summarized as the median of the performance indexes over all the sessions done on that day. Finally, the individual NF aptitude scores for each participant were defined as the mean over the daily performance of the last two days:

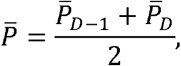

where *D* is the NF training days assigned to the participant.

### 2.4 Functional connectivity

#### Preprocessing

The resting-state fMRI data were preprocessed using SPM12 (Wellcome Trust Centre for Neuroimaging, University College London, UK) on MATLAB^®^ as described in (Ichikawa et al., 2020; Nakano et al., 2020; Yahata et al., 2016). The functional images were preprocessed with slice-timing correction, realigned to the mean image, normalized and resampled in 2 × 2 × 2 mm^3^ voxels, and smoothed with an isotropic 6-mm full-width half-maximum Gaussian kernel. The potential confounding effects (i.e., the temporal fluctuations of the white matter, the cerebrospinal fluid, and the entire brain as well as six head motion parameters) were linearly regressed out from the fMRI time series. Then a band-pass filter (0.008–0.1 Hz) was applied.

#### ROIs selection

A previous meta-analysis study reportedly found a common pattern of regional activation and deactivation during neurofeedback trials (Emmert et al., 2016). It determined 19 key brain regions of interest associated with NF. We predetermined the key regions of interest (ROIs) from Emmert et al. (ref, see Supplementary Table 1 for MNI coordinates). Of the 19 regions across the entire brain, we focused on a subset of 8 ROIs (13 regions) due to the limitations of the computational resources and for the simplicity of the interpretation focusing on the putatively relevant key brain regions. For each individual, the time-courses of the fMRI data were extracted for each of the 13 ROIs (radius = 8 mm): bilateral anterior insular cortex (AIC), bilateral dorsolateral prefrontal cortex (DLPFC), bilateral temporal parietal junction (TPJ), bilateral intraparietal sulcus (IPS), precuneus (PCn), posterior cingulate cortex (PCC), bilateral posterior insular cortex (PIC), and right superior parietal lobule (SPL). These regions belong to major functional networks and are of specific concerns to fMRI neurofeedback (salience network, executive network, attention network, default mode network, auditory network) (Emmert et al., 2016).

#### Assessing functional connectivity

Pair-wise Pearson correlations among 13 ROIs were calculated to obtain a matrix of 78 FCs for each participant. We then averaged the bilateral FCs and finally reduced to 28 FCs consisting of 8 ROIs.

### 2.5 Prediction of individual’s NF aptitude

To construct the prediction model of the individual NF aptitude scores (response variables) based on their corresponding resting-state FCs (explanatory variables) defined in the previous sections, we applied partial least squares (PLS) regression (Chen et al., 2009; Giessing et al., 2007; McIntosh et al., 2004, 1996; Wold, 1975; Yoshida et al., 2017; Ziegler et al., 2013), which is more suitable for cases with high-dimensional explanatory variables than ordinary least-square linear regression.

As possible sets of explanatory variables, we can consider in total 2^28^ = 268,435,456 combinations among the 28 FCs we assessed in the previous procedure. To extract the reasonable combinations to predict individual NF aptitudes, we adopted an exhaustive search strategy (Figure 2). In this process, we trained a PLS model for every possible combination using the discovery dataset collected at the Hiroshima University, and evaluated its predictive performance by leave-one-out cross-validation (LOOCV): One subject was used for validation data to evaluate the generalization ability of the PLS model trained by the remaining subjects; and the procedure was repeated until all the subjects were used as validation data. To cope with the high variability of individual NF aptitudes, the prediction performance was evaluated by Spearman’s correlation of true NF aptitudes and predicted scores. We applied this procedure to all possible combinations. Subsequently, we ranked the FCs in descending order based on their frequency included in the best 100 combinations in terms of prediction performance to determine the effective predictive factors for NF aptitude.

**Figure 2:**
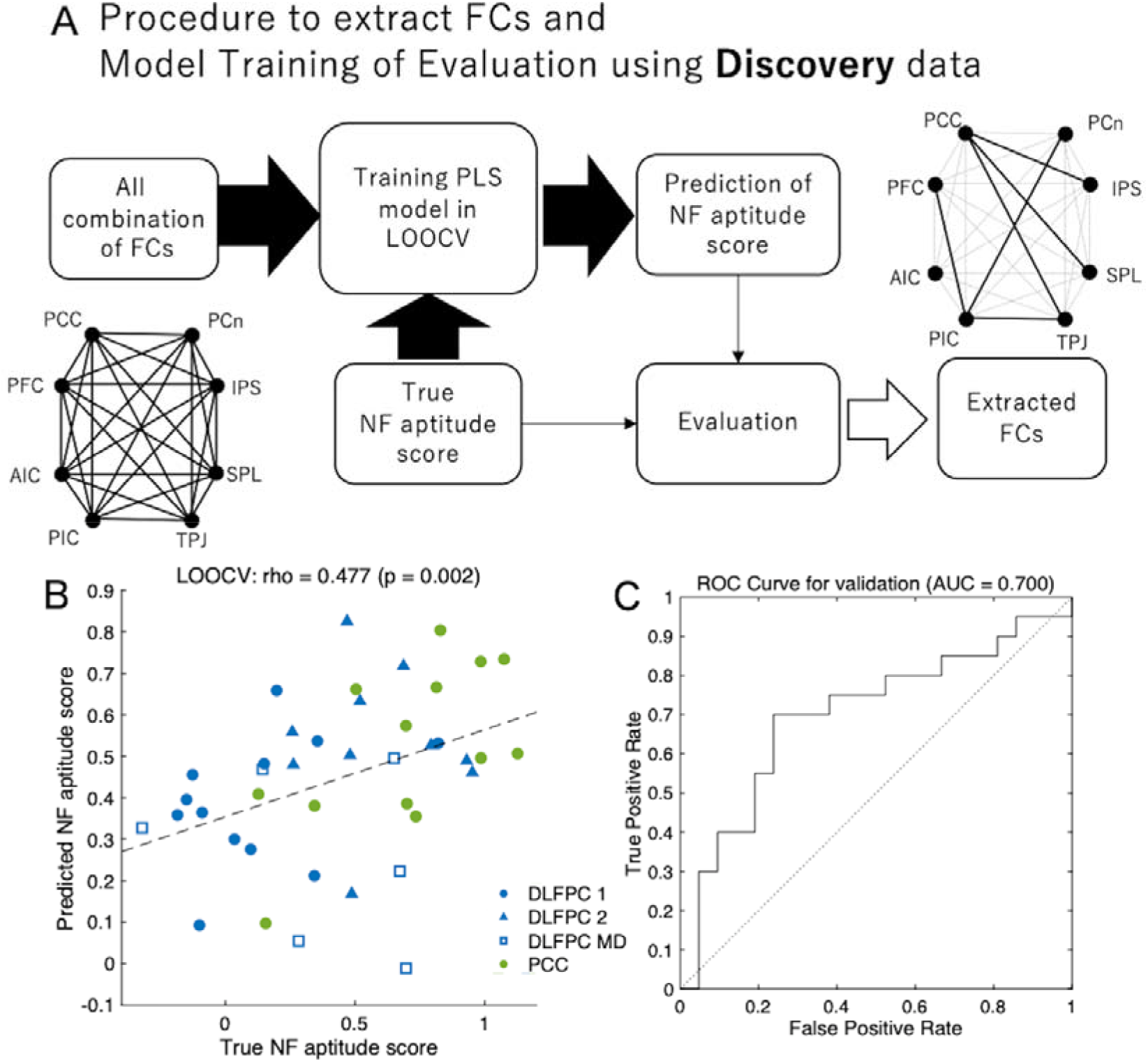
Training of the NF aptitude prediction model and extraction of the contributing functional connectivities (FCs) using the discovery data. (A) Schematic illustration of the procedures of model training and FC selection. The PLS model was used for the prediction model. All combinations connecting the eight pre-defined functional ROIs were tried one by one as the possible explanatory variables of the PLS models, and their performances were evaluated through LOOCV procedure in the discovery data. (B) NF aptitude scores predicted by the best PLS model in LOOCV of the discovery data. The scatter plot shows the true (horizontal axis) and predicted (vertical axis) NF aptitude scores validated over all runs in LOOCV. The blue markers indicate the participants using DLPFC as a target region, and green markers indicate the participants using PCC. Three different protocols in the NF targeting DLPFC were discriminated by three different marker shapes. (C) ROC curve when the best PLS model was used as binary classifier to predict whether each participant had high NF aptitude (with larger NF aptitude scores than the median in the discovery dataset) or low NF aptitude (with lower NF aptitude scores).

We also evaluated how accurately the PLS models classified the participants into high-aptitude (participants whose NF aptitude scores exceeded the median) and the low-aptitude groups (the other participants). To this end, the area under the curve of receiver operating characteristic curve, abbreviated as AUC below, were calculated in the LOOCV procedure.

Finally, we evaluated the prediction performance of the PLS models for the test dataset collected in the National Institute of Radiological Sciences, National Institutes for Quantum and Radiological Science and Technology, to demonstrate the generalization ability for the independent NF protocols. Herein, the models were trained using all the training datasets, and the prediction performance scores were identical as in the LOOCV procedure (i.e. Spearman’s correlation of the true NF aptitudes, and the predicted scores; and AUC).

## 3. Results

First, we investigated the variability of the individual NF aptitudes among the participants. The NF aptitude scores were distributed widely. Most participants (53/65) showed higher NF aptitude scores than the baseline (=0) whereas approximately 18% (12/65) of them were worse than the baseline. There were significant differences in the NF aptitude scores between the PCC-targeting and several other groups (one-way ANOVA with Scheffe post-hoc test (significance level 0.05)), but not between any other pair except PCC-targeting group (See Supplemental figure 1). Regarding the dependence on the depressive symptoms, the NF aptitude scores did not significantly differ between the MDD patients (DLPFC-MD) and the healthy control (DLPFC-2) groups. Taken together, the NF aptitude scores did not significantly depend on the depressive symptoms or the NF protocols except the PCC-targeting protocol.

Next we trained and evaluated the PLS regression model to predict the NF aptitude scores based on the resting-state FCs according to the procedure explained in Sec. 2.5. As a result of LOOCV, the PLS model with the explanatory variables, which consist of six FCs (PCC-SPL, PCC-TPJ, PCC-IPS, PIC-TPJ, PIC-DLPFC, PIC-PCn), showed the highest Spearman’s correlation between the predicted and true scores (ρ = 0.477, p =0.002, see Figure 2). When the model classified the participants into the high- or low-aptitude groups, the ACU was 0.700 (Figure 2).

We then examined whether the PLS model with the highest performance in the LOOCV could predict NF aptitude scores in the test dataset in which the target ROIs were pgACC and IPS. The predicted NF aptitude scores still considerably correlated with the true scores (ρ = 0.454, p = 0.027, AUC = 0.660; Figure 3), where the resulting regression equation was given by

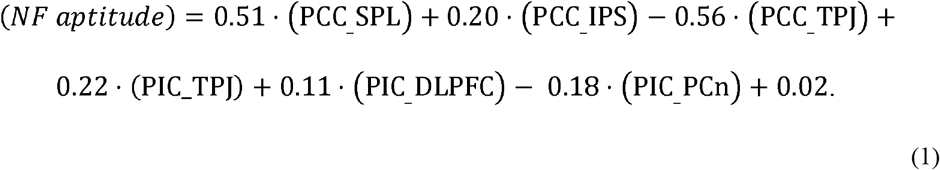

Herein, “(x_y)” denotes an FC connecting two regions of “x” and “y.”

**Figure 3:**
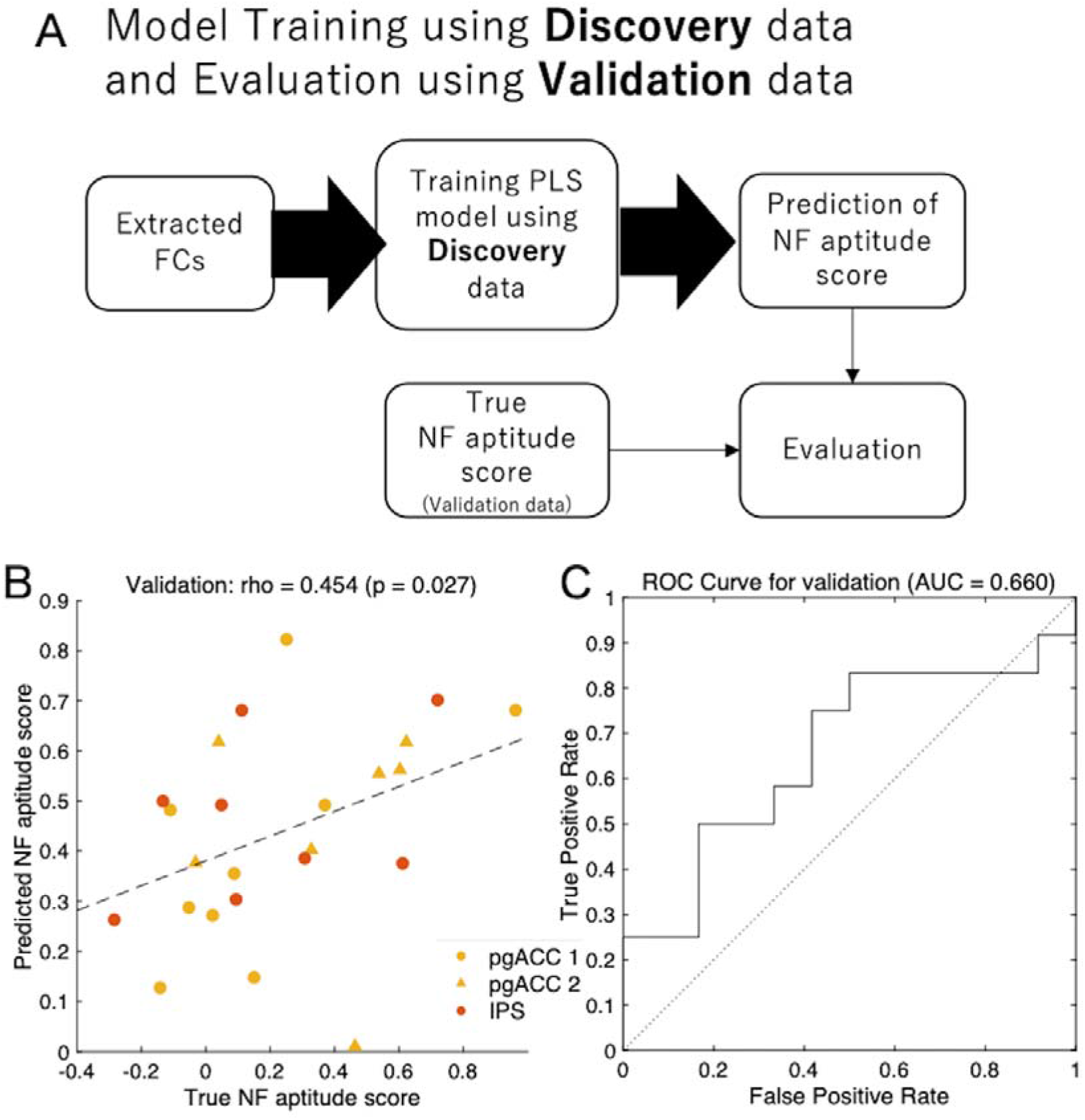
Performance evaluation of the NF aptitude prediction model scores using the test dataset. (A) Schematic illustration of the model evaluation process using the test dataset. The NF aptitude scores were predicted by the best PLS model determined in LOOCV. (B) The NF scores of the validation data were predicted using the best combination of FCs in LOOCV in the discovery data. The scatter plot shows the true (horizontal axis) and predicted (vertical axis) NF aptitude scores tested in this process. The orange markers indicate the participants using pgACC as a target region, and green markers indicate those using IPS. Two different protocols in the NF targeting pgACC are discriminated by two different marker shapes. (C) ROC curve for the test dataset when the best PLS model was used as the binary classifier to predict whether each participant had high NF aptitude (with larger NF aptitude scores than the median in the test dataset) or low NF aptitude (with lower NF aptitude scores).

**Figure 4:**
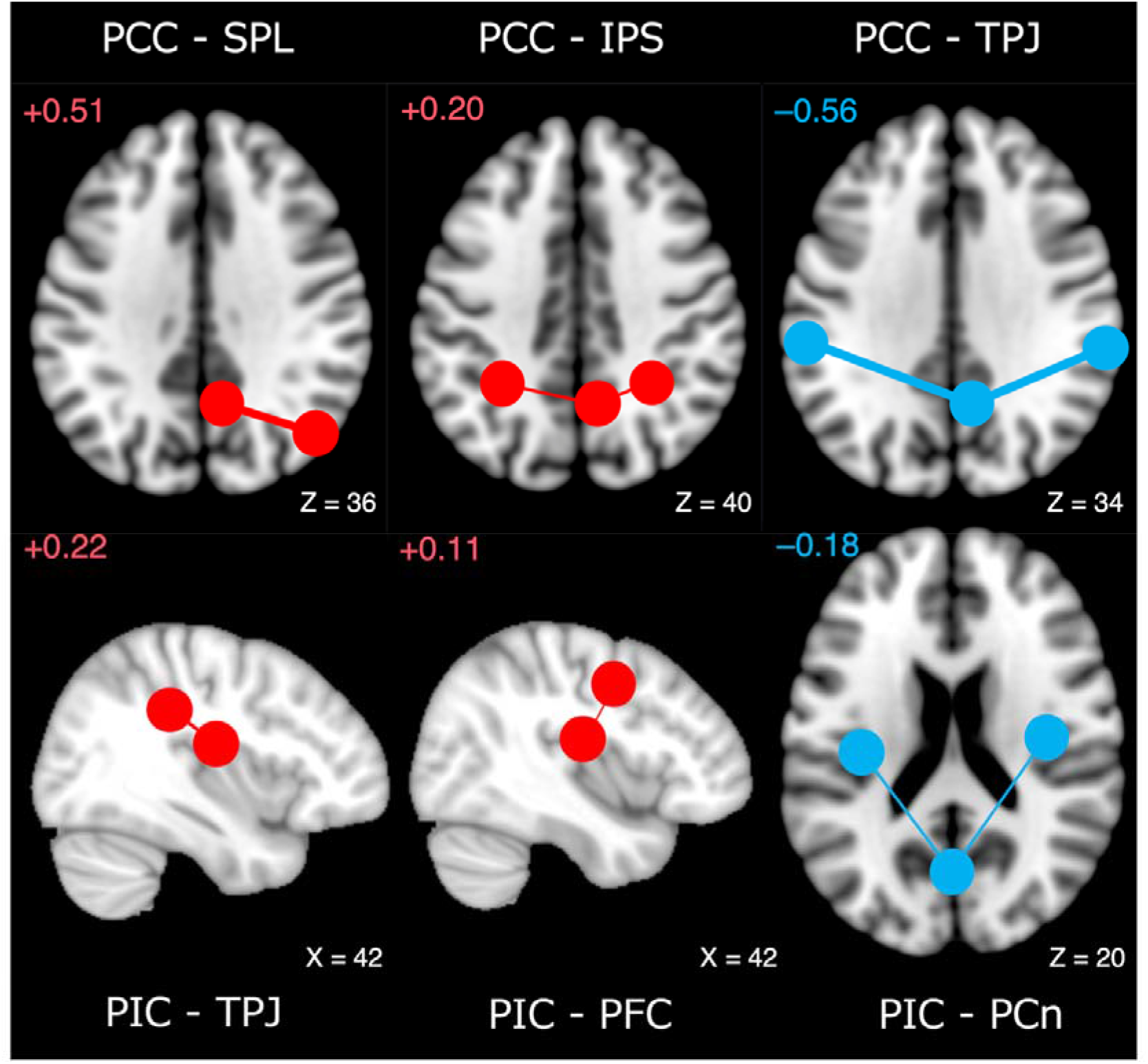
Selected functional connectivities as predictive factors for NF aptitude scores. Connectivity with positive relations with NF aptitude are colored in red and those with negative relations are colored in blue. MNI coordinates of Z or X axes are noted. Numbers in the upper left corners are regression coefficients in Equation (1). Abbreviations: posterior cingulate cortex (PCC); superior parietal lobule (SPL); intraparietal sulcus (IPS); temperoparietal junction (TPJ); posterior insular cortex (PIC); dorsolateral prefrontal cortex (PFC); precuneus (PCn).

Since it might be possible that the best PLS model was too complicated to fit the dataset, we identified FCs that were robustly selected in highly-ranked PLS models through an exhaustive search. In fact, the regression coefficients for two FCs (PCC-SPL and PCC-TPJ) were more than double as much as the others. To examine the robustness of the FCs, we checked the explanatory variables used in the best 100 PLS models and found that the above six FCs appeared more frequently than the other FCs among those models (Supplemental Figure 2). More interestingly, three FCs connected to PCC (PCC-TPJ, PCC-IPS and PCC-SPL) were always selected as the explanatory variables in the best 100 PSL models. Furthermore, the PLS model whose explanatory variables only consisted of those three FCs still showed a high prediction performance: ρ = 0.437, p =0.005 and AUC = 0.681 in the LOOCV; and ρ = 0.370, p = 0.076 and AUC = 0.632 for the test dataset (Supplemental Figure 3), which are comparable to (but not better than) the best PLS model.

## 4. Discussions

In the present study, we examined the feasibility that individual’s NF aptitudes independent of target brain regions can be predicted based on the resting-state FCs in the whole brain before NF training. The PLS model, which selected a small number of FCs as explanatory variables through a LOOCV using the training dataset, was successfully generalized to an independent test dataset. Such predictive screening is necessary for applying fMRI-NF to therapy situations. The present method demonstrated a preferable property in that the predictive screening could be achieved only by low-burden resting state fMRI scanning.

In the best PLS model, PCC and PIC were functional hubs connecting six ROIs that were selected as explanatory variables. Among them, it was notable that the FCs between PCC and other ROIs were robustly selected in the highly-ranked PLS models through an exhaustive search. PCC, which is a part of the default mode network (Buckner et al., 2008), plays an important role in self-oriented processing such as retrieving autobiographical memory or self-referential processing (Leech and Sharp, 2014). PIC is related to interoceptive information processing (Craig, 2002). These functions may be related to NF aptitude.

On the other hand, regions considered to be directly involved in control during NF, such as DLPFC and AIC (Emmert et al., 2016), were not selected as hubs. This result suggests that the active control of those regions during tasks is important, but not reflected in connectivity in the task-free resting-state.

For comparisons with existing related studies, (Scheinost et al., 2014) and (Haugg et al., 2020b) predicted the appropriateness of NFs from brain indices of resting-state FCs and brain activity in the localizer tasks, respectively. In both studies, the brain indices used for prediction corresponded to each NF target. On the other hand, our present study demonstrated a feasibility that the NF aptitude of multiple target regions can be predicted by the common brain indices. For example, even if we consider only depression, various candidate targets (e.g. amygdala, insula, DLPFC) have been suggested (Linden, 2014). Thus, the prediction model presented here might be used more widely than the existing prediction models for specific NF targets. As a future prospect, it might be more clinically valuable if a precise prediction of which types and targets of NF treatment are suitable for an individual could be made from resting-state MRI alone.

A limitation of this study is that the number of the ROIs was restricted to enable an exhaustive search for the selection of explanatory variables. It is possible that better models could be created from whole brain networks with a larger number of nodes. Methodologically, the development of automatic variable selection alternatives to exhaustive searches is a future issue. In addition, more sites and within-subject replications are also needed to reinforce the generalization ability.

Features other than resting functional images may also be useful in the predictions. Examination using indices such as gray matter volume, cortical thickness, white matter fibers, or neurotransmitters may be useful not only to construct better predictive models but also to explore the neuroscientific basis of adaptation to NF training.

## Acknowledgements

This research was supported by the Strategic Research Program for Brain Sciences (Integrated Research on Depression, Dementia and Development Disorders) from the Japan Agency for Medical Research and Development, AMED, Grant Numbers JP20dm0107093, JP20dm0107094, and JP20dm0107096. This work was also partially supported by JSPS KAKENHI Grant Numbers JP20K06874 and JP20H00625.

## Data and code availability statements

The original data for this study will not be made publicly available due to the involvement of patient data. It can be obtained upon request to the Department of Psychiatry and Neurosciences, Hiroshima University, Japan (SY). IRB imposing these restrictions on our data is Ethical Committee for Epidemiology of Hiroshima University (contact: Shoji Karatsu). Additional contact information is available from: https://www.hiroshima-u.ac.jp/en/pharm/contact.

## Credit authorship contribution statement

**Takashi Nakano:** Methodology, Software, Investigation, Writing - original Draft, Writing - Review & Editing. **Masahiro Takamura:** Conceptualization, Methodology, Resources, Writing - original Draft, Writing - Review & Editing. **Haruki Nishimura:** Methodology, Resources, Writing - original Draft, Writing - Review & Editing. **Maro Machizawa:** Methodology, Resources, Writing - original Draft, Writing - Review & Editing. **Naho Ichikawa:** Resources. **Atsuo Yoshino:** Resources. **Go Okada:** Resources. **Yasumasa Okamoto:** Conceptualization, Supervision. **Shigeto Yamawaki:** Supervision, Funding acquisition. **Makiko Yamada:** Methodology, Resources, Writing - original Draft, Writing - Review & Editing. **Tetsuya Suhara:** Supervision, Funding acquisition. **Junichiro Yoshimoto:** Conceptualization, Methodology, Software, Investigation, Writing - original Draft, Writing - Review & Editing, Funding acquisition.

## Supplementary materials

**Supplemental Table 1:** The ROIs selected in this study and their MNI coordinates. The ROI names, their abbreviations, and their original ROI names used in the paper by (Emmert et al., 2016). We used the identical coordinates as in their paper.

| ROI name | Abbreviation | Name in (Emmert et al., 2016) | MNI Coordinates |
| --- | --- | --- | --- |
|  |  |  | (x, y, z) |
| anterior insular cortex | AIC | anterior insular cortex | 32, 26, 4 (R) |
|  |  |  | -36, 20, -2 (L) |
| dorsolateral prefrontal cortex | PFC | dorsolateral prefrontal cortex | 42, 0, 42 (R) |
|  |  |  | -34, -4, 40 (L) |
| temporal parietal junction | TPJ | Temporo-parietal | 62, -34, 34 (R) |
|  |  |  | -58, -32, 32 (L) |
| intraparietal Sulcus | IPS | Parietal | 30, -48, 40 (R) |
|  |  |  | -30, -48, 38 (L) |
| posterior cingulate cortex | PCC | posterior cingulate cortex | 8, -56, 38 |
| precuneus | PCn | precuneus | 0, -68, 24 |
| posterior insular cortex | PIC | Temporal transverse | -36, -20, 16 (R) |
|  |  |  | 38, -14, 18 (L) |
| superior parietal lobule | SPL | Parietal R | 46, -68, 36 |

**Supplemental Figure 1:**
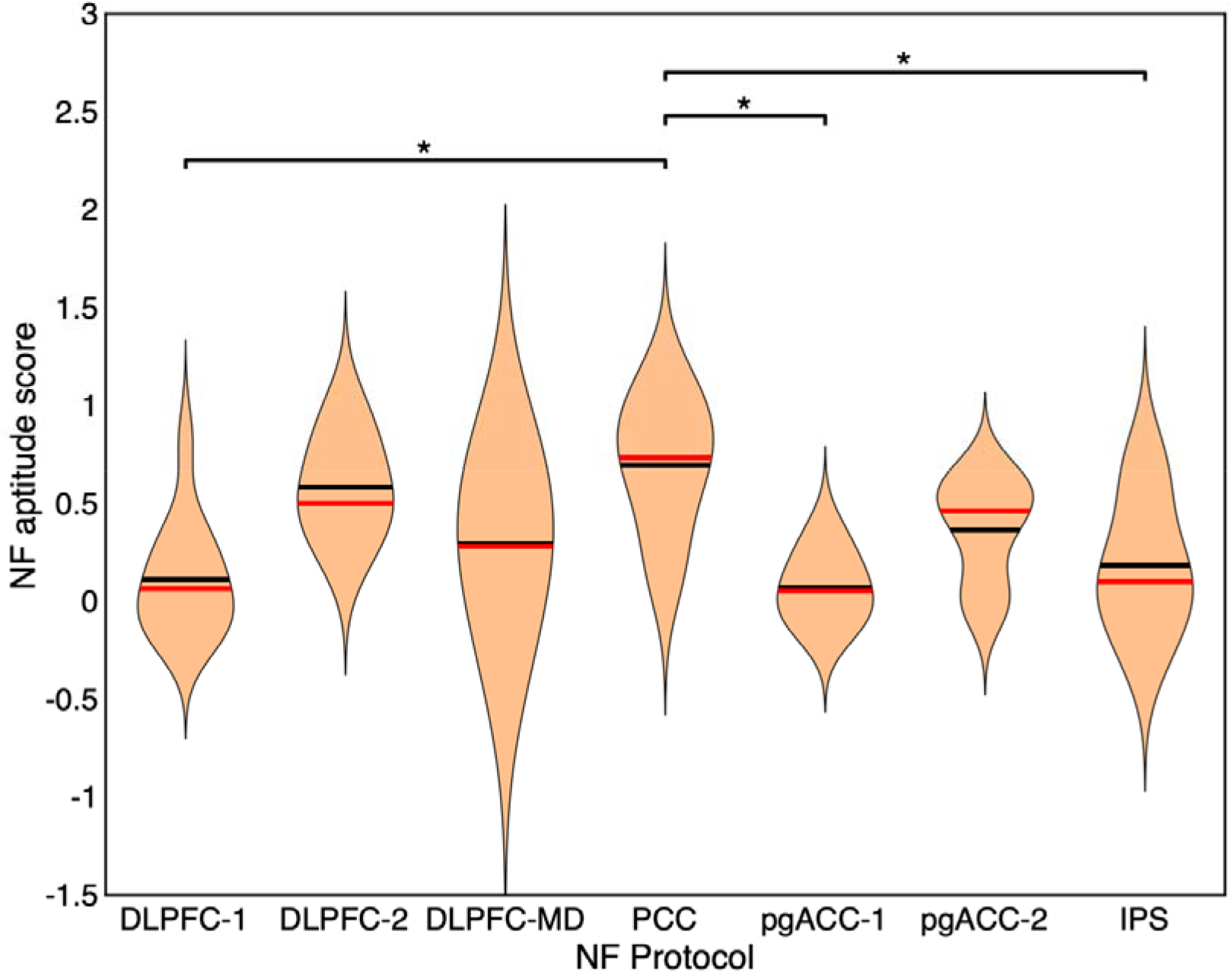
Distribution of NF aptitude scores for each NF group. The asterisks (*) indicate the significant differences (significance level: 0.05, one-way ANOVA and Scheffe post-hoc test) between groups.

**Supplemental Figure 2:**
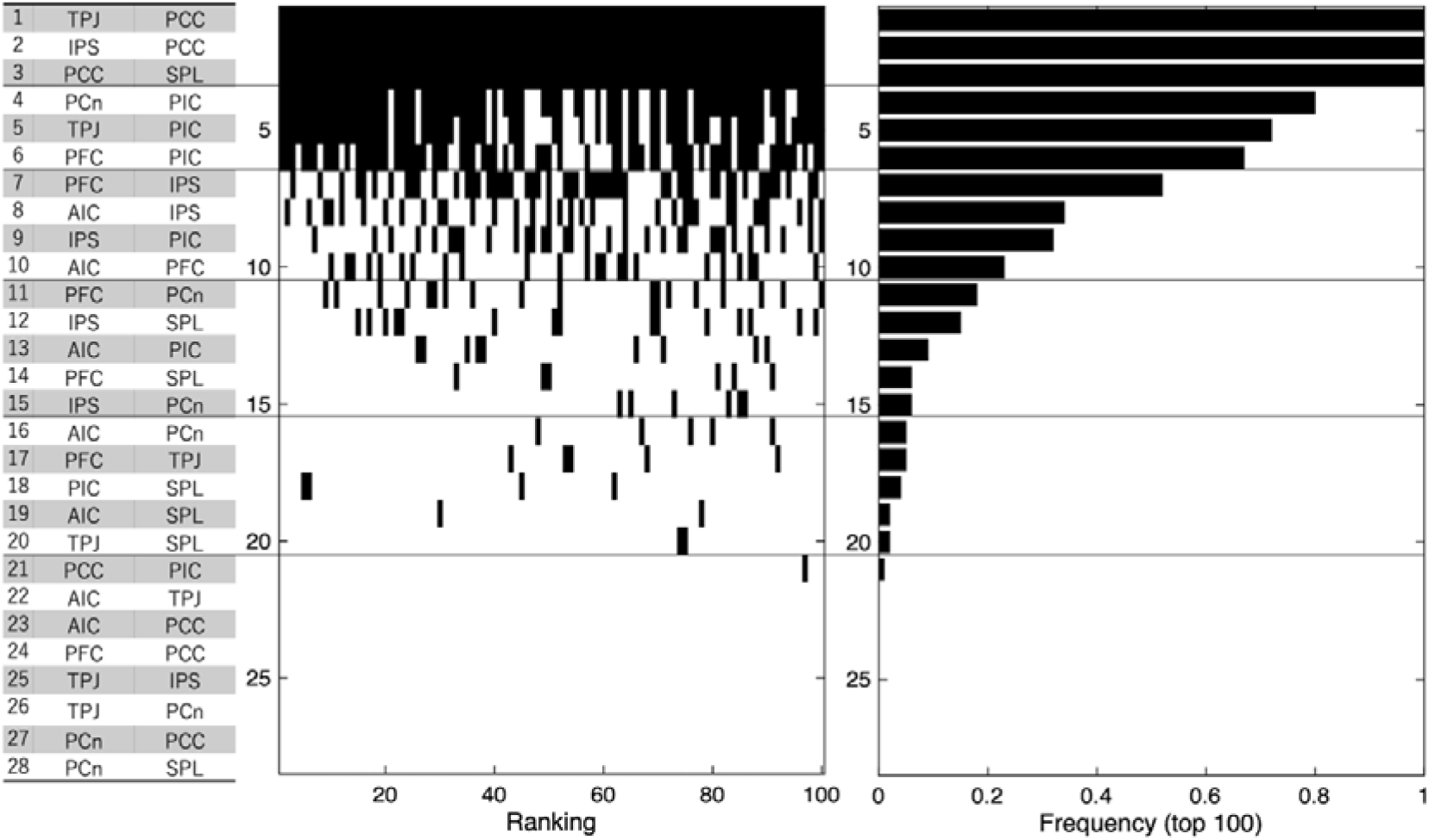
List of functional connectivities (FCs) sorted in the descending order of the frequency included in the best 100 prediction models. (Left) The list of FCs. (Center) The indicators whether the FC for each row was used (black) or not (white). The horizontal axis denotes the rank of the PLS model, based on the prediction performance in the LOOCV. (Right) The frequency of FCs used in the best 100 PSL models.

**Supplemental Figure 3:**
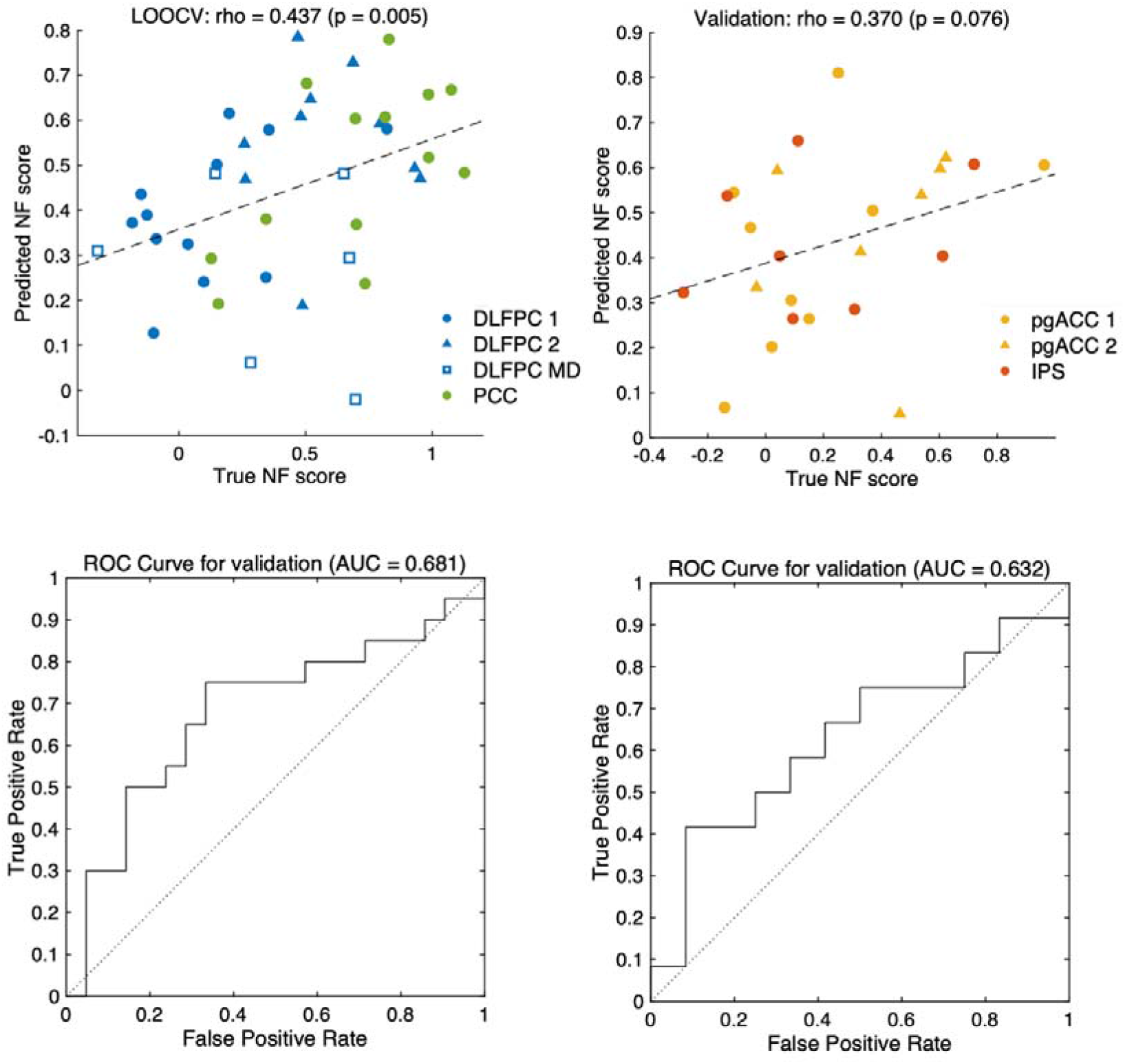
Prediction performance of the PLS model with only 3 FCs connecting PCC as the explanatory variables. The performance was evaluated by the LOOCV using the discovery dataset (Left) and generalization to the independent test dataset (Right). Upper panels: The scatter plot of the true (horizontal axis) and predicted (vertical axis). Lower panels: ROC curve when the best PLS model was used for the binary classifier to predict whether each participant had high NF aptitude (with larger NF aptitude score than the median in the corresponding dataset) and low NF aptitude (with lower NF aptitude score).

## References

Alegria, A.A., Wulff, M., Brinson, H., Barker, G.J., Norman, L.J., Brandeis, D., Stahl, D., David, A.S., Taylor, E., Giampietro, V., Rubia, K., 2017. Real-time fMRI neurofeedback in adolescents with attention deficit hyperactivity disorder. Hum. Brain Mapp. 38, 3190–3209. https://doi.org/10.1002/hbm.23584

Buckner, R.L., Andrews-Hanna, J.R., Schacter, D.L., 2008. The Brain’s Default Network. Ann. N. Y. Acad. Sci. 1124, 1–38. https://doi.org/10.1196/annals.1440.011

Chen, K., Reiman, E.M., Huan, Z., Caselli, R.J., Bandy, D., Ayutyanont, N., Alexander, G.E., 2009. Linking functional and structural brain images with multivariate network analyses: A novel application of the partial least square method. Neuroimage 47, 602–610. https://doi.org/10.1016/j.neuroimage.2009.04.053

Craig, A.D., 2002. How do you feel? Interoception: the sense of the physiological condition of the body. Nat. Rev. Neurosci. 3, 655–666. https://doi.org/10.1038/nrn894

Emmert, K., Kopel, R., Sulzer, J., Brühl, A.B., Berman, B.D., Linden, D.E.J., Horovitz, S.G., Breimhorst, M., Caria, A., Frank, S., Johnston, S., Long, Z., Paret, C., Robineau, F., Veit, R., Bartsch, A., Beckmann, C.F., Van De Ville, D., Haller, S., 2016. Meta-analysis of real-time fMRI neurofeedback studies using individual participant data: How is brain regulation mediated? Neuroimage 124, 806–812. https://doi.org/10.1016/j.neuroimage.2015.09.042

Gerin, M.I., Fichtenholtz, H., Roy, A., Walsh, C.J., Krystal, J.H., Southwick, S., Hampson, M., 2016. Real-Time fMRI Neurofeedback with War Veterans with Chronic PTSD: A Feasibility Study. Front. Psychiatry 7, 1–11. https://doi.org/10.3389/fpsyt.2016.00111

Giessing, C., Fink, G.R., Rösler, F., Thiel, C.M., 2007. fMRI Data Predict Individual Differences of Behavioral Effects of Nicotine: A Partial Least Square Analysis. J. Cogn. Neurosci. 19, 658–670. https://doi.org/10.1162/jocn.2007.19.4.658

Haugg, A., Renz, F.M., Nicholson, A.A., Lor, C., Götzendorfer, S.J., Sladky, R., Skouras, S., McDonald, A., Craddock, C., Hellrung, L., Kirschner, M., Herdener, M., Koush, Y., Papoutsi, M., Keynan, J., Hendler, T., Kadosh, K.C., Zich, C., Kohl, S.H., Hallschmid, M., MacInnes, J., Adcock, A., Dickerson, K., Chen, N.-K., Young, K., Bodurka, J., Marxen, M., Yao, S., Becker, B., Auer, T., Schweizer, R., Pamplona, G., Lanius, R.A., Emmert, K., Haller, S., Ville, D. Van De, Kim, D.-Y., Lee, J.-H., Marins, T., Fukuda, M., Sorger, B., Kamp, T., Liew, S.-L., Veit, R., Spetter, M., Weiskopf, N., Scharnowski, F., Steyrl, D., Van De Ville, D., Kim, D.-Y., Lee, J.-H., Marins, T., Fukuda, M., Sorger, B., Kamp, T., Liew, S.-L., Veit, R., Spetter, M., Weiskopf, N., Scharnowski, F., Steyrl, D., Ville, D. Van De, Kim, D.-Y., Lee, J.-H., Marins, T., Fukuda, M., Sorger, B., Kamp, T., Liew, S.-L., Veit, R., Spetter, M., Weiskopf, N., Scharnowski, F., Steyrl, D., 2020a. Determinants of Real-Time fMRI Neurofeedback Performance and Improvement – a Machine Learning Mega-Analysis. bioRxiv 2020.10.21.349118. https://doi.org/10.1101/2020.10.21.349118

Haugg, A., Sladky, R., Skouras, S., McDonald, A., Craddock, C., Kirschner, M., Herdener, M., Koush, Y., Papoutsi, M., Keynan, J.N., Hendler, T., Cohen Kadosh, K., Zich, C., MacInnes, J., Adcock, R.A., Dickerson, K., Chen, N.K., Young, K., Bodurka, J., Yao, S., Becker, B., Auer, T., Schweizer, R., Pamplona, G., Emmert, K., Haller, S., Van De Ville, D., Blefari, M.L., Kim, D.Y., Lee, J.H., Marins, T., Fukuda, M., Sorger, B., Kamp, T., Liew, S.L., Veit, R., Spetter, M., Weiskopf, N., Scharnowski, F., 2020b. Can we predict real-time fMRI neurofeedback learning success from pretraining brain activity? Hum. Brain Mapp. 41, 3839–3854. https://doi.org/10.1002/hbm.25089

Ichikawa, N., Lisi, G., Yahata, N., Okada, G., Takamura, M., Hashimoto, R. ichiro, Yamada, T., Yamada, M., Suhara, T., Moriguchi, S., Mimura, M., Yoshihara, Y., Takahashi, H., Kasai, K., Kato, N., Yamawaki, S., Seymour, B., Kawato, M., Morimoto, J., Okamoto, Y., 2020. Primary functional brain connections associated with melancholic major depressive disorder and modulation by antidepressants. Sci. Rep. 10, 1–12. https://doi.org/10.1038/s41598-020-60527-z

Ito, T., Yokokawa, K., Yahata, N., Isato, A., Suhara, T., Yamada, M., 2017. Neural basis of negativity bias in the perception of ambiguous facial expression. Sci. Rep. 7, 420. https://doi.org/10.1038/s41598-017-00502-3

Kadosh, K.C., Staunton, G., 2019. A systematic review of the psychological factors that influence neurofeedback learning outcomes. Neuroimage 185, 545–555. https://doi.org/10.1016/j.neuroimage.2018.10.021

Leech, R., Sharp, D.J., 2014. The role of the posterior cingulate cortex in cognition and disease. Brain 137, 12–32. https://doi.org/10.1093/brain/awt162

Linden, D.E.J., 2014. Neurofeedback and networks of depression. Dialogues Clin. Neurosci. 16, 103–112. https://doi.org/10.31887/DCNS.2014.16.1/dlinden

McIntosh, A.R., Bookstein, F.L., Haxby, J. V, Grady, C.L., 1996. Spatial pattern analysis of functional brain images using partial least squares. Neuroimage 3, 143–157. https://doi.org/10.1006/nimg.1996.0016

McIntosh, A.R., Chau, W.K., Protzner, A.B., 2004. Spatiotemporal analysis of event-related fMRI data using partial least squares. Neuroimage 23, 764–775. https://doi.org/10.1016/j.neuroimage.2004.05.018

Mehler, D.M.A., Sokunbi, M.O., Habes, I., Barawi, K., Subramanian, L., Range, M., Evans, J., Hood, K., Lührs, M., Keedwell, P., Goebel, R., Linden, D.E.J., 2018. Targeting the affective brain—a randomized controlled trial of real-time fMRI neurofeedback in patients with depression. Neuropsychopharmacology 43, 2578–2585. https://doi.org/10.1038/s41386-018-0126-5

Nakano, T., Takamura, M., Ichikawa, N., Okada, G., Okamoto, Y., Yamada, M., Suhara, T., Yamawaki, S., Yoshimoto, J., 2020. Enhancing Multi-Center Generalization of Machine Learning-Based Depression Diagnosis From Resting-State fMRI. Front. Psychiatry 11, 1–10. https://doi.org/10.3389/fpsyt.2020.00400

Paret, C., Kluetsch, R., Zaehringer, J., Ruf, M., Demirakca, T., Bohus, M., Ende, G., Schmahl, C., 2016. Alterations of amygdala-prefrontal connectivity with real-time fMRI neurofeedback in BPD patients. Soc. Cogn. Affect. Neurosci. 11, 952–960. https://doi.org/10.1093/scan/nsw016

Ruiz, S., Lee, S., Soekadar, S.R., Caria, A., Veit, R., Kircher, T., Birbaumer, N., Sitaram, R., 2013. Acquired self-control of insula cortex modulates emotion recognition and brain network connectivity in schizophrenia. Hum. Brain Mapp. 34, 200–212. https://doi.org/10.1002/hbm.21427

Scheinost, D., Stoica, T., Saksa, J., Papademetris, X., Constable, R.T., Pittenger, C., Hampson, M., 2013. Orbitofrontal cortex neurofeedback produces lasting changes in contamination anxiety and resting-state connectivity. Transl. Psychiatry 3, e250–e250. https://doi.org/10.1038/tp.2013.24

Scheinost, D., Stoica, T., Wasylink, S., Gruner, P., Saksa, J., Pittenger, C., Hampson, M., 2014. Resting state functional connectivity predicts neurofeedback response. Front. Behav. Neurosci. 8, 1–7. https://doi.org/10.3389/fnbeh.2014.00338

Sepulveda, P., Sitaram, R., Rana, M., Montalba, C., Tejos, C., Ruiz, S., 2016. How feedback, motor imagery, and reward influence brain self-regulation using real-time fMRI. Hum. Brain Mapp. 37, 3153–3171. https://doi.org/10.1002/hbm.23228

Sherwood, M.S., Kane, J.H., Weisend, M.P., Parker, J.G., 2016. Enhanced control of dorsolateral prefrontal cortex neurophysiology with real-time functional magnetic resonance imaging (rt-fMRI) neurofeedback training and working memory practice. Neuroimage 124, 214–223. https://doi.org/10.1016/j.neuroimage.2015.08.074

Sitaram, R., Ros, T., Stoeckel, L., Haller, S., Scharnowski, F., Lewis-Peacock, J., Weiskopf, N., Blefari, M.L., Rana, M., Oblak, E., Birbaumer, N., Sulzer, J., 2017. Closed-loop brain training: the science of neurofeedback. Nat. Rev. Neurosci. 18, 86–100. https://doi.org/10.1038/nrn.2016.164

Stoeckel, L.E., Garrison, K.A., Ghosh, S.S., Wighton, P., Hanlon, C.A., Gilman, J.M., Greer, S., Turk-Browne, N.B., DeBettencourt, M.T., Scheinost, D., Craddock, C., Thompson, T., Calderon, V., Bauer, C.C., George, M., Breiter, H.C., Whitfield-Gabrieli, S., Gabrieli, J.D., LaConte, S.M., Hirshberg, L., Brewer, J.A., Hampson, M., Van Der Kouwe, A., Mackey, S., Evins, A.E., 2014. Optimizing real time fMRI neurofeedback for therapeutic discovery and development. NeuroImage Clin. 5, 245–255. https://doi.org/10.1016/j.nicl.2014.07.002

Subramanian, L., Hindle, J. V., Johnston, S., Roberts, M. V., Husain, M., Goebe, R., Linden, D., 2011. Real-time functional magnetic resonance imaging neurofeedback for treatment of Parkinson’s disease. J. Neurosci. 31, 16309–16317. https://doi.org/10.1523/JNEUROSCI.3498-11.2011

Takamura, M., Okamoto, Y., Shibasaki, C., Yoshino, A., Okada, G., Ichikawa, N., Yamawaki, S., 2020 Antidepressive effect of left dorsolateral prefrontal cortex neurofeedback in patients with major depressive disorder: A preliminary report. J. Affect. Disord. 271, 224–227. https://doi.org/10.1016/j.jad.2020.03.080

Thibault, R.T., MacPherson, A., Lifshitz, M., Roth, R.R., Raz, A., 2018. Neurofeedback with fMRI: A critical systematic review. Neuroimage 172, 786–807. https://doi.org/10.1016/j.neuroimage.2017.12.071

Wold, H., 1975. Path Models with Latent Variables: The NIPALS Approach, in: Quantitative Sociology. Elsevier, pp. 307–357. https://doi.org/10.1016/B978-0-12-103950-9.50017-4

Yahata, N., Morimoto, J., Hashimoto, R., Lisi, G., Shibata, K., Kawakubo, Y., Kuwabara, H., Kuroda, M., Yamada, T., Megumi, F., Imamizu, H., Náñez Sr, J.E., Takahashi, H., Okamoto, Y., Kasai, K., Kato, N., Sasaki, Y., Watanabe, T., Kawato, M., 2016. A small number of abnormal brain connections predicts adult autism spectrum disorder. Nat. Commun. 7, 11254. https://doi.org/10.1038/ncomms11254

Yamashita, M., Yoshihara, Y., Hashimoto, R., Yahata, N., Ichikawa, N., Sakai, Y., Yamada, T., Matsukawa, N., Okada, G., Tanaka, S.C., Kasai, K., Kato, N., Okamoto, Y., Seymour, B., Takahashi, H., Kawato, M., Imamizu, H., 2018. A prediction model of working memory across health and psychiatric disease using whole-brain functional connectivity. Elife 7, 1–26. https://doi.org/10.7554/eLife.38844

Yoshida, K., Shimizu, Y., Yoshimoto, J., Takamura, M., Okada, G., Okamoto, Y., Yamawaki, S., Doya, K., 2017. Prediction of clinical depression scores and detection of changes in whole-brain using resting-state functional MRI data with partial least squares regression. PLoS One 12, e0179638. https://doi.org/10.1371/journal.pone.0179638

Young, K.D., Misaki, M., Harmer, C.J., Victor, T., Zotev, V., Phillips, R., Siegle, G.J., Drevets, W.C., Bodurka, J., 2017a. Real-Time Functional Magnetic Resonance Imaging Amygdala Neurofeedback Changes Positive Information Processing in Major Depressive Disorder. Biol. Psychiatry 82, 578–586. https://doi.org/10.1016/j.biopsych.2017.03.013

Young, K.D., Siegle, G.J., Zotev, V., Phillips, R., Misaki, M., Yuan, H., Drevets, W.C., Bodurka, J., 2017b. Randomized Clinical Trial of Real-Time fMRI Amygdala Neurofeedback for Major Depressive Disorder: Effects on Symptoms and Autobiographical Memory Recall. Am. J. Psychiatry 174, 748–755. https://doi.org/10.1176/appi.ajp.2017.16060637

Ziegler, G., Dahnke, R., Winkler, A.D., Gaser, C., 2013. Partial least squares correlation of multivariate cognitive abilities and local brain structure in children and adolescents. Neuroimage 82, 284–294. https://doi.org/10.1016/j.neuroimage.2013.05.088

Zilverstand, A., Sorger, B., Sarkheil, P., Goebel, R., 2015. fMRI neurofeedback facilitates anxiety regulation in females with spider phobia. Front. Behav. Neurosci. 9, 1–12. https://doi.org/10.3389/fnbeh.2015.00148

Zweerings, J., Hummel, B., Keller, M., Zvyagintsev, M., Schneider, F., Klasen, M., Mathiak, K., 2019. Neurofeedback of core language network nodes modulates connectivity with the default-mode network: A double-blind fMRI neurofeedback study on auditory verbal hallucinations. Neuroimage 189, 533–542. https://doi.org/10.1016/j.neuroimage.2019.01.058

